# aPKC-mediated displacement and actomyosin-mediated retention polarize Miranda in *Drosophila* neuroblasts

**DOI:** 10.1101/148213

**Authors:** Matthew Hannaford, Anne Ramat, Nicolas Loyer, Jens Januschke

## Abstract

Cell fate generation can rely on the unequal distribution of molecules during progenitor cell division in the nervous system of vertebrates and invertebrates. Here we address asymmetric fate determinant localization in the developing *Drosophila* nervous system, focussing on the control of asymmetric Miranda distribution in larval neuroblasts. We used live imaging of neuroblast polarity reporters at endogenous levels of expression to address Miranda localization during the cell cycle. We reveal that the regulation and dynamics of cortical association of Miranda in interphase and mitosis are different. In interphase Miranda binds directly to the plasma membrane. At the onset of mitosis, Miranda is phosphorylated by aPKC and displaced from the PM. After nuclear envelope breakdown asymmetric localization of Miranda requires actomyosin activity. Therefore, Miranda phosphorylation by aPKC and differential binding to the actomyosin network are required at distinct phases of the cell cycle to polarize fate determinant localization.

## INTRODUCTION

The development of the central nervous system depends on asymmetric cell divisions for the balanced production of progenitor as well as differentiating cells. During vertebrate and invertebrate neurogenesis, cell fates can be established through the unequal inheritance of cortical domains or fate determinants during asymmetric progenitor divisions (Knoblich 2008; Doe 2008; Marthiens & ffrench-Constant 2009; Alexandre et al. 2010).

An important step in asymmetric cell division is the establishment of a polarity axis. Asymmetrically dividing *Drosophila* neuroblasts (NBs) establish an axis of polarity at the onset of mitosis. Like in many other polarized cells, this depends on the activity of the Par complex (Goldstein & Macara 2007). As NBs enter prophase, the Par complex including Par3/Bazooka (Baz), aPKC and Par-6 assembles at the apical NB pole, which drives the localization of fate determinants to the opposite NB pole establishing the apico basal polarity axis (Wodarz et al. 1999; Wodarz et al. 2000; Rolls et al. 2003; Petronczki & Knoblich 2001; Betschinger et al. 2003; Homem & Knoblich 2012; Prehoda 2009).

Upon NB division, the basally localized fate determinants segregate to the daughter cell that commits to differentiation. For this to happen correctly, two adapter proteins, Partner of Numb (Pon, Lu et al. 1998) and Miranda (Mira, Ikeshima-Kataoka et al. 1997; Shen et al. 1997), are required. While Pon localizes the Notch signaling regulator Numb (Uemura et al. 1989; Lu et al. 1998), Mira localizes the homeobox transcription factor Prospero (Pros) and the translational repressor Brat to the basal NB cortex in mitosis (Ikeshima-Kataoka et al. 1997; Betschinger et al. 2006; C.-Y. Lee et al. 2006). In the absence of Mira, fate determination is impaired and tumor-like growth can occur in larval NB lineages (Ikeshima-Kataoka et al. 1997; Caussinus & Gonzalez 2005).

How polarized fate determinant localization in mitotic NBs is achieved is a long-standing question and despite being a well-studied process, this mechanism is not fully understood. In embryonic NBs, Mira localization requires actin (Shen et al. 1998) and myosin activity since mutation in the myosin regulatory light chain *spaghetti squash (sqh)* (Barros et al. 2003) or the Myosin VI gene *jaguar* (Petritsch et al. 2003) to lead to Mira localization defects. Furthermore, in embryos injected with the Rho kinase (ROCK) inhibitor Y-27632, Mira does not achieve a polarized distribution, which was rescued by the expression of a phospho-mimetic *sqh* allele. This led to the proposal that Myosin II plays a critical role in Mira localization and aPKC affects Mira localization indirectly through regulating Myosin II (Barros et al. 2003).

However, it was later shown that Y-27632 can inhibit aPKC directly and that Mira is a substrate of aPKC (Atwood & Prehoda 2009; Wirtz-Peitz et al. 2008). In fact many aPKC substrates including Numb and Mira contain a basic and hydrophobic (BH) motif that can be phosphorylated by aPKC. Upon phosphorylation the substrates are no longer able to directly bind phospholipids of the plasma membrane (PM, Bailey & Prehoda 2015; Smith et al. 2007; Dong et al. 2015). Asymmetric Mira localization in mitotic NBs can in principle be explained by keeping the activity of aPKC restricted to the apical pole (Atwood & Prehoda 2009). However, is less clear what the contribution of the actomyosin network to polarized Mira localization is.

Intriguingly, Mira localizes uniformly to the cortex of larval brain NBs in interphase (Sousa-Nunes et al. 2009) before becoming basally restricted in mitosis. The function and regulation of interphase Mira are unknown. In embryonic NBs in interphase, Mira and its cargo Pros localize to the apical cortex (Spana & Doe 1995) and Pros is found in the nucleus of interphase *mira* mutant NBs (Matsuzaki et al. 1998). Given that the levels of nuclear Pros in NBs are important for the regulation of entry and exit from quiescence and NB differentiation (Lai & Doe 2014), interphase Mira localization might be important in this context.

To reveal the regulation of interphase Mira, we set out to determine differences and similarities in the parameters of Mira binding in interphase and mitosis and the transition between the two different localizations. Using fluorescent reporters of the Par complex and Mira at endogenous levels of expression, we reveal that cortical Mira localization is differently controlled in interphase, where it uniformly binds to the PM and after nuclear envelope breakdown (NEB), when actomyosin activity becomes important.

## RESULTS

### Uniform Miranda is cleared from the cortex during prophase and Miranda reappears asymmetrically after NEB

We first confirmed in larval NBs that Mira localizes uniformly to the cortex in interphase (**Figure 1A**) and co-localizes with its cargo Pros in interphase and mitosis (**Figure 1 supplement 1**). We further stained *mira* mutant NBs for Mira and Pros. In these cells, Pros is found in the nucleus in interphase (**Figure 1 supplement 1**). These results are consistent with a role of Mira in regulation Pros localization not only in mitosis (Ikeshima-Kataoka et al. 1997), but also potentially during interphase. We therefore sought to address the regulation of cortical Mira in interphase and mitosis and the transition between these localizations.

**Figure 1.**
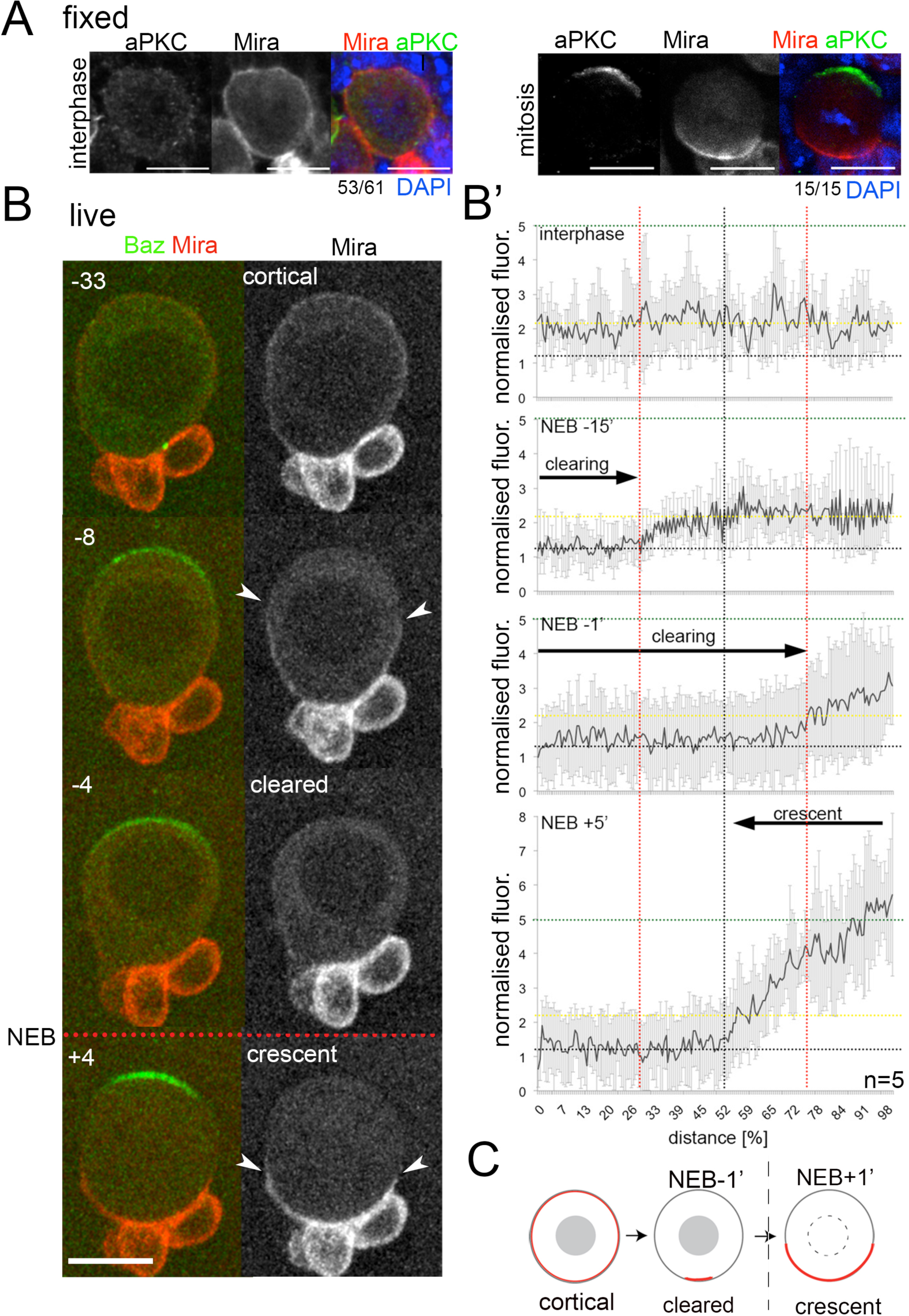
Miranda is cleared from the cortex before localizing in a basal crescent in mitosis. (**A**) Larval brain NBs fixed and stained as labeled at the indicated cell cycle stage. (**B**) Selected frames from **Movie S1**. NB in primary cell culture expressing Baz::GFP (green) and Mira::mCherry (red) in the transition from interphase to mitosis. Arrowheads point at Mira being cleared (-8) and at basal Mira crescent (+4). (**B**’) Quantification of cortical Mira::mCherry signal plotting the fluorescence intensities from the apical to the basal pole computationally straightening (Kocsis et al. 1991) the cortices of 5 NBs against the distance in %. Fluorescence was background subtracted and normalized to background subtracted cytoplasmic signal (1, dotted line). Cortical signal (yellow dotted line) and signal after NEB (green dotted line). Error bars, standard deviation. (**C**) Schematic of Mira localization. *BAC{mira::mcherry-MS2}* was the source of Mira::mCherry. Scale bar 10µm. Time stamp: minutes.

To accurately monitor *in vivo* the dynamics of this transition, we used a BAC construct in which Mira was tagged at its C-terminus with mCherry (Ramat et al. 2017; **Figure 1 supplement 2**). This tagged Mira recapitulated uniform cortical localization in interphase (**Figure 1B**, **-**33 to NEB) and polarized localization to the basal pole in mitosis (**Figure 1B**, +4), with a 2.5-fold increase in intensity (n=5, **Figure 1B’**). This transition occurred in two distinct steps. Firstly, during prophase, after Mira was rapidly excluded from the apical pole where Baz (**Movie S1**) or aPKC (**Movie S2**) started localizing (**Figure 1B -**8 to NEB, **B’** -15). Thereafter, Mira was progressively cleared from most of the rest of the cortex in an apical-to-basal direction (**Figure 1B, B’** -4, -1, respectively to NEB). Secondly, following NEB, Mira reappeared at the cortex in a basal crescent after NEB (**Figure 1B,B’** +4, +5 respectively). We further recapitulated these steps using overexpression of GFP::Mira and by antibody staining of endogenous, non-tagged Mira. The Mira antibody revealed colcemid insensitive cortical localization in interphase, which was lost in *mira* mutant NBs (**Figure 1 supplement 3**). Thus, endogenous Mira also localizes to the cortex independently of microtubules. In conclusion, Mira transitions from a uniform to a polarized localization by first being cleared form most of the cortex during prophase, and only upon NEB reappearing in a basal crescent (**Figure 1C**).

### Actomyosin is required for the establishment and maintenance of Miranda crescents after NEB

In embryonic NBs, Mira localization is sensitive to F-actin disruption in mitosis (Shen et al. 1998). We reasoned that this could be due to defective Mira clearance in prophase and/or defective Mira reappearance at the basal cortex after NEB. We tested this by disrupting the actin network of larval NBs with Latrunculin A (LatA). Despite efficiently disrupting F-actin (**Figure 2 supplement**) and causing cytokinesis failure (**Figure 2A,** 3:02, related to **Movie S3**), LatA treatment affected neither uniform cortical localization of Mira in interphase (**Figure 2A,** 2:06) nor its clearance during prophase (**Figure 2A,** 2:21). In contrast, Mira failed to relocalize to a basal crescent following NEB in LatA-treated NBs (**Figure 2A,** 2:33). We next tested whether, in addition to crescent establishment, F-Actin controlled crescents maintenance. To this end, we arrested NBs with colcemid in metaphase (at which point Mira crescents are established), after which we treated them with LatA. Again, LatA caused Mira to relocalize to the cytoplasm (**Figure 2A**). Importantly, this relocalization is unlikely to be due to Mira being “swept off” the cortex by cortical aPKC, which redistributes from an apical crescent to the entire cortex following LatA treatment (**Figure 2 supplement**). This is evident since unlike the gradual apical-to-basal clearance of Mira during prophase (**Figure 1B**), basal Mira crescents fall off the cortex homogenously upon LatA treatment (**Figure 2 supplement**). Furthermore, Mira falls off the cortex 2.8±1min (n=13) before aPKC (**Movie S4**) or Baz become detectable at the basal cortex (**Figure 2 supplement**). Therefore, an intact actin network is required to establish and maintain asymmetric Mira localization in mitotic larval NBs.

**Figure 2.**
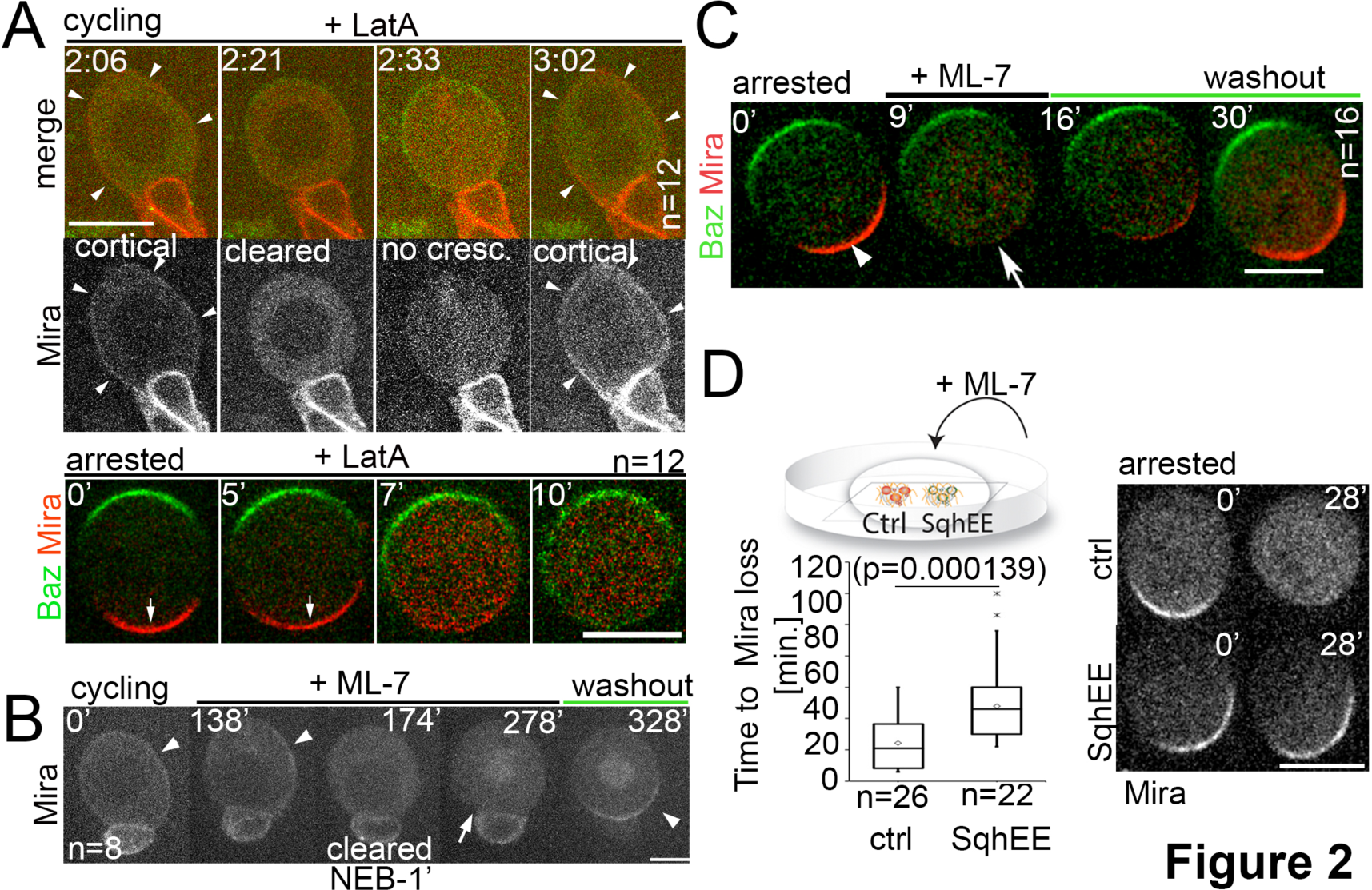
Differential response of Mira localization in interphase and mitosis to disruption of the actin cytoskeleton. (**A**) Stills from **Movie S3**. LatA was added to a cycling NB in primary cell culture expressing Baz::GFP (green) and Mira::mCherry (red). Arrowheads point at cortical Mira after culturing ~1h with LatA (2:06). At 1 min to NEB, Mira::mCherry is cleared from the cortex (2:21). Mira forms no crescent in the next mitosis (2:33), but after cytokinesis fails (note bi-nucleated cell at 3:02), Mira is recruited to the cortex (arrowheads). Bottom panels: Colcemid arrested NBs expressing Baz::GFP and Mira::mCherry. 5µM LatA was added prior to imaging at 15sec. intervals. Mira crescents (arrows) are lost upon LatA treatment. (**B**) Cycling NB in primary cell culture expressing Mira::mCherry, that remains cortical upon ML-7 addition (15µM; interphase: 0’ and 138’, arrowheads), is cleared 1minute prior to NEB (174’, does not form a crescent after NEB (278’, arrow), but accumulates on the spindle (seen in cross section). After ML-7 washout, a basal Mira::mCherry crescent recovers (arrowhead, 328’). (**C**) Related to **Movie S5.** Colcemid arrested NB in primary cell culture expressing Baz::GFP (green) and Mira::mCherry (red). After addition of 20µM ML-7 Mira (arrowhead, 0′)becomes cytoplasmic (arrow, +9’), but upon ML-7 washout a Mira crescent recovers. (**D**) The effect of 20µM ML-7 can be quenched by overexpressing a phospho-mimetic form Sqh (Sqh^EE^). Colcemid arrested NBs (ctrl: Mira::mCherry: SqhEE: Mira::mCherry co-expressing SqhEE by worniuGal4). Ctrl and SqhEE were co-cultured and ML-7 added (related to **Movie S6**). Quantification of the time required to cause Mira::mCherry to become cytoplasmic shown on the left. Two-tailed ttest for independent means revealed significance. *BAC{mira::mcherry-MS2}* was the source of Mira::mCherry. Scale bar: 10µm.

Myosin activity has been proposed to be required for Mira localization (Petritsch et al. 2003; Barros et al. 2003). We tested next which step of Mira localization involved Myosin. Myosin motor activity relies on the phosphoregulation of myosin regulatory light chain, encoded by the *sqh* gene in *Drosophila* (Jordan & Karess 1997), which we disrupted by applying ML-7, a specific inhibitor of the myosin light chain kinase (MLCK, Bain et al. 2003). As with LatA, the ML-7 treatment in cycling neuroblasts affected neither uniform cortical localization of Mira in interphase (**Figure 2B,** 0’) nor its clearance during prophase (**Figure 2B,** 138-278’) but resulted in failure to establish a basal crescent after NEB, which was restored upon drug washout (**Figure 2B**, 328’). In colcemid-arrested NBs, the ML-7 treatment also resulted in Mira redistributing from the basal crescent to the cytoplasm, which was restored upon ML-7 washout. Furthermore, unlike LatA treatment, ML-7 did not cause the Par complex to redistribute to the entire cortex (**Figure 2C, Movie S5**). Finally, as in (Das & Storey 2014), we demonstrated the specificity of the effect of ML-7 by counteracting its effect with a phosphomimetic version of Myosin regulatory light chain, Sqh. Overexpressing a phosphomimetic version of Sqh (Sqh^EE^, (Winter et al. 2001) significantly delayed loss of cortical Mira after ML-7 addition in colcemid arrested NBs (**Figure 2D, Movie S6**).

In conclusion, the contribution of F-Actin (Shen et al. 1998) and Myosin (Petritsch et al. 2003; Barros et al. 2003) to asymmetric Mira localization is linked to the relocalization of Mira and its maintenance at the basal cortex following NEB, but not to the control of uniform cortical localization in interphase or to clearance of Mira during prophase.

### Defective clearance of interphase cortical Miranda by aPKC results in the persistence of uniform Miranda in mitosis

NBs mutant for aPKC also fail to asymmetrically localize Mira in mitosis (Rolls et al. 2003), prompting us to investigate which aspect of Mira localization involved aPKC in larval NBs. Like in controls, Mira localized uniformly to the cortex of *apkc^k06403^* (Wodarz et al. 2000) mutant NBs in interphase. However, it did not clear from the cortex during prophase and instead remained uniformly localized throughout mitosis (**Movie S7**, **Figure 3A**). Additionally, expressing a constitutively active form of aPKC, aPKC^ΔN^, resulted in the loss of uniform cortical Mira binding in interphase (**Figure 3 supplement**). These results show that during prophase, aPKC negatively regulates the uniform cortical localization of Mira observed in interphase. They also suggest that the failure to asymmetrically localize Mira in mitosis in aPKC mutants is due to defective clearance in prophase resulting in the abnormal persistence of uniform cortical Mira throughout mitosis.

**Figure 3.**
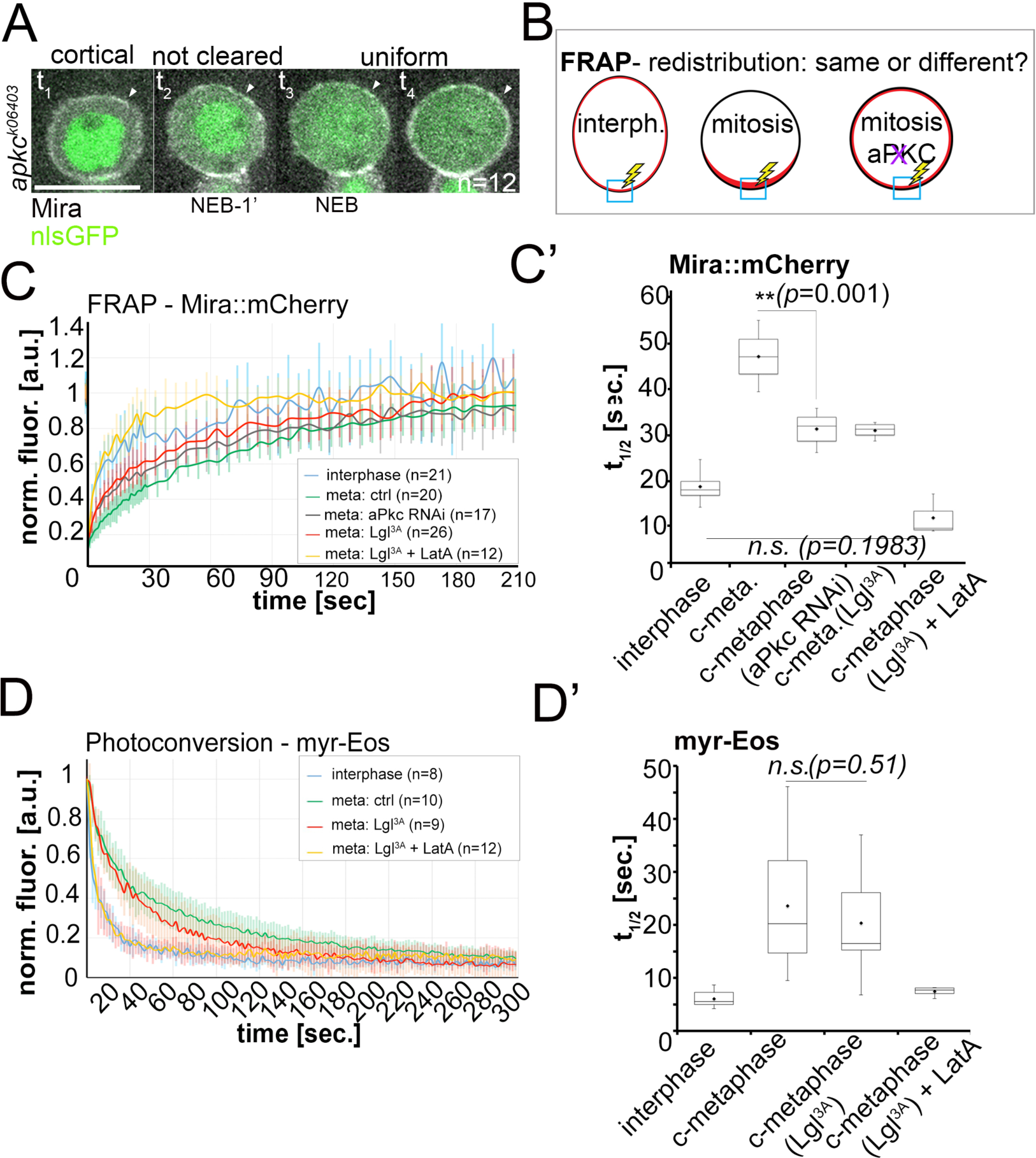
Lateral diffusion and cytoplasmic exchange of cortical Miranda are different in control and aPKC impaired mitotic NBs. (**A**) Stills from **Movie S7** of an *apkc^k06403^* mutant NB (MARCM clone labeled with nlsGFP, green) expressing Mira::mCherry (grey). Mira is cortical in interphase, as the NB enters mitosis and after NEB (arrowheads, t_1_ – t_4_). (**B**) Conditions analyzed by FRAP. (**C**) Fluorescence redistribution curves of cortical Mira::mCherry at the indicated conditions. (**C’**) Estimates of t_1/2_ [sec.] for cortical Mira::mCherry under the indicated conditions derived from curve fitting (Rapsomaniki et al. 2012). (**D**) Photo-conversion experiment monitoring loss of myr-EOS converted signal over time. (**D’**) Estimates of t_1/2_ [sec.] for cortical Mira::mCherry under the indicated conditions from curve fitting. Overexpression was driven by worniu-Gal4. p values: two-tailed ttest for independent means. Scale bar: 10µm.

We reasoned that, if this were the case, Mira localization in interphase should have the same characteristics as Mira localization in metaphase upon loss of function of aPKC. Consistent with this view, unlike Mira localization in mitosis, but similar to Mira localization in interphase (**Figure 2A**), Mira localization to the cortex in mitosis was insensitive to LatA treatment in NBs depleted for aPKC (**Figure 3 supplement**). To corroborate these results we used fluorescence redistribution after photo-bleaching (FRAP) to analyze the lateral diffusion/cytoplasmic exchange dynamics of Mira (**Figure 3B**).

In controls, Mira redistributed about three times faster in interphase compared to mitosis (**Figure 3C, C’**), suggesting that Mira dynamics are indeed a relevant characteristic to analyze cell cycle-dependent control of Mira localization. Yet, in apparent contradiction with our proposition, while Mira redistribution became faster when aPKC was knocked down by RNAi (or Lgl^3A^ overexpression, known to inhibit aPKC, Betschinger et al. 2003), these conditions did not result in Mira redistribution in mitosis becoming as fast as in interphase (**Figure 3C, C’**). However, changes in the actin network caused by progression through the cell cycle (Ramanathan et al. 2015) can influence dynamics of membrane-associated proteins in general (Heinemann et al. 2013). This is also the case in NBs, as a photo-convertible membrane-associated reporter that attaches to the entire NB PM via a myristoylation signal (myr-Eos) showed similar slowing of dynamics in mitosis compared to interphase (~four-fold, **Figure 3D, D’**).

Thus, this general cell cycle-driven change of dynamics may account for the difference between Mira redistribution in interphase and in mitosis in aPKC-impaired NBs. We tested this by cancelling this general change by treating mitotic NBs with LatA, which resulted as expected in myr-Eos dynamics falling into ranges similar to interphase (**Figure 3D, D’**). In aPKC-impaired NBs, the same LatA treatment resulted in Mira redistribution rates becoming as fast as in interphase (Figure 3C, C’), suggesting that aPKC loss of function indeed leads to the persistence of the same dynamics as in interphase (**Figure 3C, C’**). Importantly, myr-Eos dynamics were not sensitive to Lgl^3A^ overexpression alone, arguing against the possibility that aPKC impairment causes general changes in the cortex that could explain accelerated redistribution of Mira in this condition.

In conclusion, instead of being cleared, Mira persists throughout mitosis with the same actin-insensitive uniform localization and the same dynamics as in interphase in *apkc* mutant NBs.

### Miranda binds uniformly to the plasma membrane in interphase

In *Drosophila* S2 cells Mira binds directly to phospholipids of the PM via its BH motif and the ability to bind the PM is abolished upon phosphorylation of this motif by aPKC (Bailey & Prehoda, 2015). Our results, that Mira cortical localization in interphase is actin-independent and that its clearance in prophase is aPKC dependent, suggest that in interphase Mira is retained uniformly at the PM by its BH motif.

Five aPKC phosphorylation sites have been identified in Mira, only one of which Serine 96 (S96) directly resides in the BH motif. To investigate its influence on dynamic Mira localization in neuroblasts, we used CrispR to generate mCherry-tagged versions of Mira control (S96, ctrl, able to rescue embryonic lethality; see **Figure 1 supplement 2**), a phosphomutant version (S96A, homozygous embryonic lethal), a phosphomimetic version (S96D, homozygous embryonic lethal) - shown *in vitro* to reduce phospholipid binding and Mira recruitment to the PM when overexpressed in S2 cells (Bailey & Prehoda 2015) - and a complete deletion of the BH motif (ΔBH, homozygous embryonic lethal).

The control, S96, localized uniformly to the cortex in interphase, was cleared from the cortex in prophase and reappeared as a basal crescent after NEB (**Figure 4A, Movie S8**). In contrast, while S96A localized uniformly to the cortex in interphase, it was not cleared at the onset of prophase when it presented a transient apical enrichment, perhaps explainable by abnormally stable interactions with apically localized aPKC. After NEB, S96A, localized however uniformly to the cortex (**Figure 4A**, **Movie S9**), which occurred independently of F-actin (**Figure 4B**). Furthermore, S96D did not localize to the cortex in interphase and instead accumulated predominantly on cortical microtubules, as evidenced by its relocalization to the cytoplasm upon colcemid treatment (**Figure 4B**). Nonetheless, S96D always localized asymmetrically at the cortex following NEB, despite at reduced levels (**Figure 4A**, **Movie S10**).

**Figure 4.**
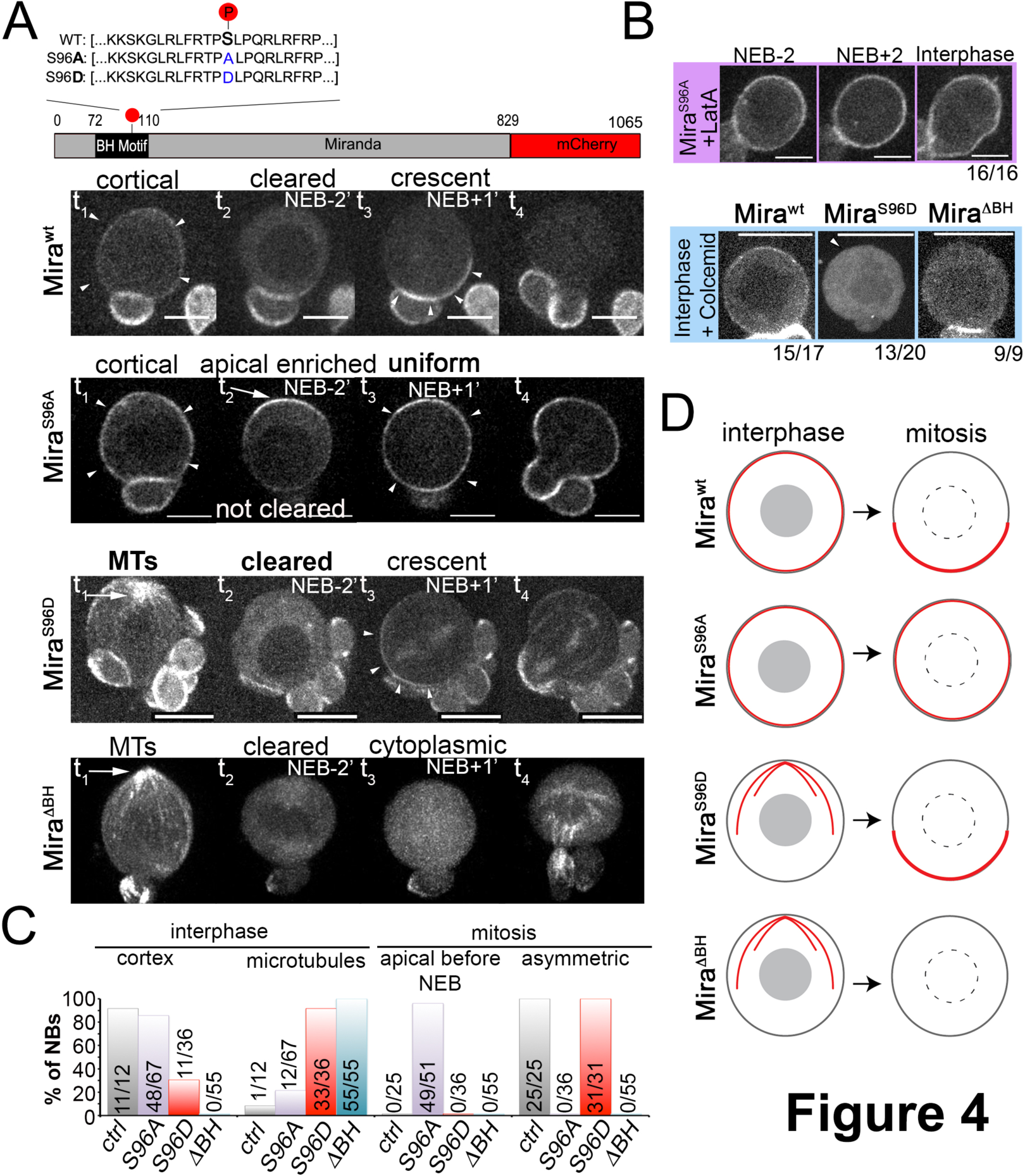
Miranda binds to the plasma membrane in interphase NBs via its BH motif. (**A**) Schematic indicating the different Mira alleles used. Mira::mCherry localizes cortically uniform in interphase (arrowheads t_1_), is cleared from the cortex shortly before NEB and forms a crescent (arrowheads t_3_) thereafter that is inherited by daughter cells (related to **Movie S8**). The phosphomutant S96A is uniformly cortical in interphase, accumulates apically shortly before NEB (arrow, t_2_), is uniformly cortical after NEB (arrowheads t_3_) and in telophase (t_4_, related to **Movie S9**). The phosphomimetic S96D localizes to cortical microtubules in interphase (arrow t_1_), is cleared from the cortex before NEB and asymmetric after NEB (arrowheads t_3_) and segregates to daughter cells (related to **Movie S10**). Deletion of the BH motif leads to cortical microtubule localization in interphase (arrow t1), cytoplasmic localization before and after NEB and reappearance on microtubules around cytokinesis (related to **Movie S11**). (**B**) Neuroblasts expressing the indicated Mira alleles were treated with 1µM LatA or 50µM colcemid for 60min. Cortical localization of S96A is insensitive to LatA treatment. Below: While the Control remains cortical, S96D and ΔBH become cytoplasmic upon colcemid treatment. (**C**) Frequency of indicated localization of the different Mira mutants. (**D**) Schematic of the localization of the different Mira alleles. Scale bar: 10µm.

Finally, deletion of the BH motif abolished both uniform cortical localization in interphase and asymmetric cortical localization after NEB (**Figure 4A**, **Movie S11,** see **Figure 4C** for quantification and **Figure 4D** for summary of the localization of the different Mira mutants in interphase and mitosis). Therefore, our findings are consistent with Mira being bound via its BH motif to the phospholipids of the PM in interphase, and that phosphorylation of S96 by aPKC in prophase disrupts this interaction.

### Mira crescent size is affected by a Y-27632-sensitive mechanism that operates before NEB

So far, our results demonstrate that in interphase Mira binds uniformly the PM, from which it is cleared by aPKC at the onset of prophase. However, upon NEB Mira localization requires actomyosin. This raises the question whether aPKC contributes to Mira asymmetric localization after NEB. Temporally controlled inactivation can be achieved with temperature sensitive (ts) alleles or small molecule inhibition. We find that the available ts allele of aPKC (Guilgur et al. 2012) is hypomorphic already at permissive temperature resulting in Mira localization defects (not shown). Therefore, we made use of the non-specific effects of the ROCK inhibitor Y-27632, which inhibits aPKC with an IC_50_ of 10µM (Atwood & Prehoda 2009). When we added 50µM of Y-27632 to already polarized NBs, after 1h in the presence of the drug, we did not detect any significant changes in Mira or aPKC crescent size (n=22, **Figure 5A,** and **Figure 5 supplement**).

**Figure 5.**
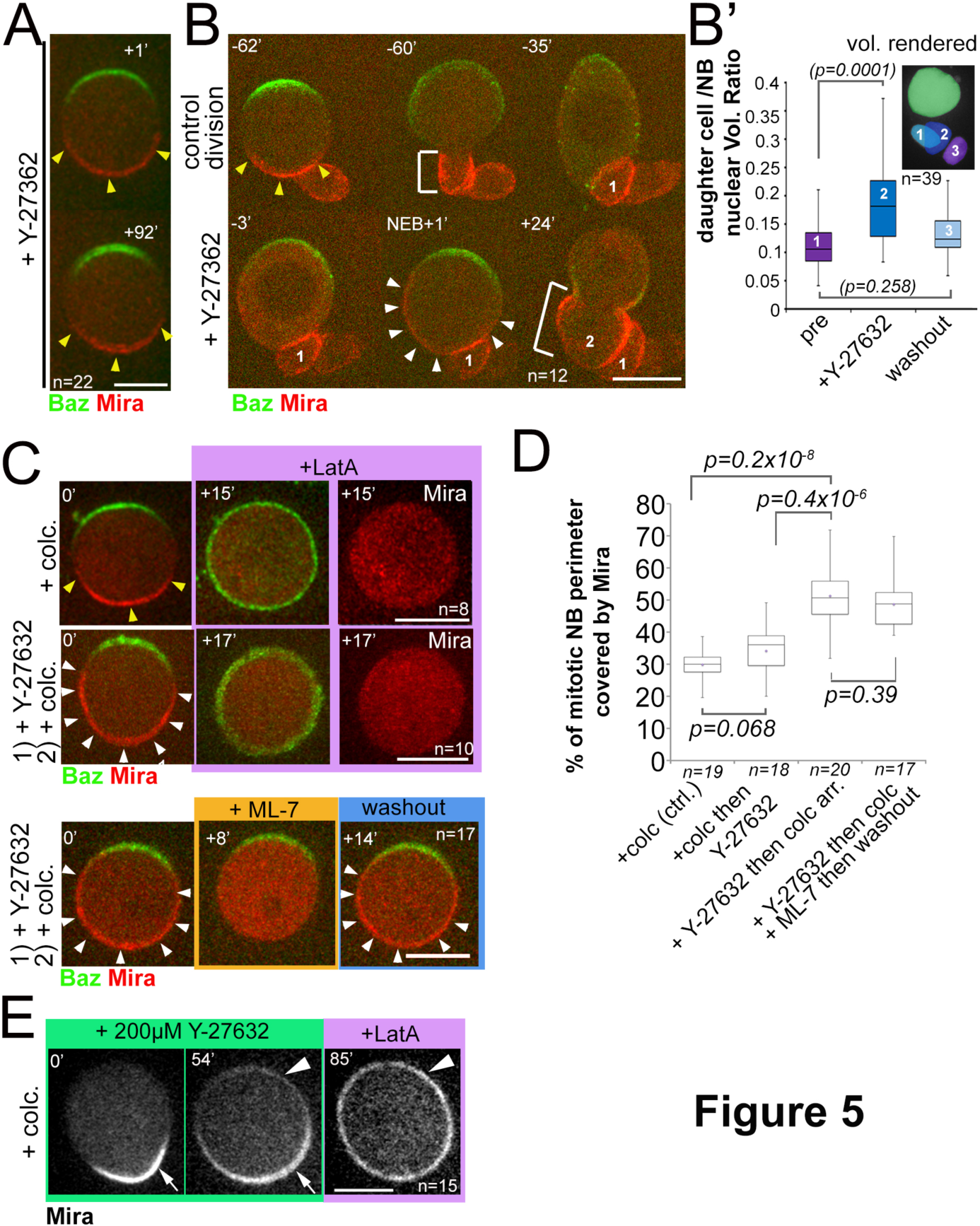
Mira crescent size is affected by a Y-27632-sensitive mechanism that operates before NEB. (**A**) Culturing colcemid arrested NBs in 50µM Y-27632 did not alter Mira crescent size (yellow arrowheads). (**B**) NBs polarizing in the presence of 25µM Y-27632 show enlarged Mira crescents. Control division (-62’ to -35’) with normally sized Mira crescent and daughter cell size (-60’; yellow arrowheads, bracket, respectively). Dividing in the presence of Y-27632 (-3, NEB+1) leads to an enlarged Mira crescent (NEB+1, white arrowheads) and enlarged daughter cell size (+24’, brackets, 2). (**B’**) Plot of the ratio of daughter cell to NB nuclei as a measure for the effect of Y-27632 on daughter cell size. NBs expressing NLSGFP were imaged by DIC to follow daughter cell birth order during three consecutive divisions [1) pre-treatment; 2) div. in the presence of 25µM Y-27632; 3) div. after drug washout]. Then a high-resolution z-stack of nlsGFP was recorded, and the nuclear volumes rendered and calculated using IMARIS to plot their ratio. *p* values: Dunn’s test. (**C**) NBs were allowed to polarize in the absence (*upper row*) or presence of 25µM Y-27632 (middle and lower row) followed by colcemid arrest. *upper row:* Control NB with normal Mira crescent (yellow arrowheads) was depolarized by 1µM LatA. Mira was displaced into the cytoplasm. *middle row:* adding 1µM LatA leads to displacement of the enlarged Mira crescent (yellow arrowheads) in the cytoplasm. *Lower row:* adding 20µM ML-7 drives Mira into the cytoplasm (+8’). Upon ML-7 washout, Mira recovered to an enlarged crescent (+14’, white arrowheads). (**D**) Quantification of Mira crescent size in the aforementioned experiments (unpaired ttest). (**E**) Colcemid arrested NBs were treated with 200µM Y-276322. Mira remains asymmetric even after 56min in the drug. LatA addition (5µM) abolishes that asymmetric bias and Mira is uniformly distributed on the membrane. *Time* stamp: min. Labels as indicated. *BAC{mira::mcherry-MS2}* was the source of Mira::mCherry. Scale bar: 10µm.

In striking contrast, when we added half that dose to cycling NBs, while aPKC crescent size was comparable to controls (**Fig 5 supplement**) Mira crescents formed, but were significantly enlarged. As a consequence daughter cell size was also increased (n=12, **Figure 5B, B**’. Intriguingly, such enlarged crescents were LatA and ML-7 sensitive (**Figure 5C**, see **Figure 5D** for Mira crescent size quantification), both indicators that enlarged Mira crescents were not due to lack of phosphorylation of Mira by aPKC. However, when we titrated the Y-27632 concentration to induce uniform cortical Mira when added to cycling NBs, Mira became uniformly cortical and was insensitive to LatA treatment in the presence of 200µM Y-27632, suggesting that this concentration can lead to inhibition of Mira phosphorylation by aPKC (**Movie S12**, n=25 and not shown).

We then treated colcemid arrested NBs with 200µM Y-27632. It took on average 52±11min (n=15) until faint Mira signal became detectable apically. Nevertheless, Mira distribution remained always biased towards the basal pole in the presence of Y-27632, an asymmetry that was lost only when we additionally added LatA, which caused Mira to become uniformly distributed on the PM (**Figure 5E**, **Movie 13**, n=15). Therefore, 200µM Y-27632 when added to colcemid arrested NBs, may have an effect on Mira phosphorylation by aPKC, causing ectopic accumulation of Mira apically. However, given the non-specific nature of Y-27632 especially at high concentrations, a role for aPKC cannot be precisely determined in this way.

Nonetheless, when added at lower concentrations (25µM) before NEB, Y-27632 has an effect on Mira crescent size that is unlikely to be caused by failure of aPKC to phosphorylate Mira.

## Discussion

The establishment of different cell fates in the developing nervous system can rely on asymmetric distribution of molecules. In this study, we reinvestigated *in vivo* the precise contributions of aPKC and actomyosin to the localization of the cell fate determinant adapter Mira during the cell cycle of *Drosophila* larval NBs. Using live cell imaging we reveal a stepwise contribution of aPKC and actomyosin to Mira localization. First, Mira was cleared from most of the cortex during prophase; second, Mira came back to the cortex as a basal crescent following NEB (**Figure 1B-C**). We then proceeded to assess the relative contributions of aPKC and actomyosin to these steps. Our study reveals that the central elements of two models proposed to explain Mira asymmetry, requiring actomyosin (Barros et al. 2003) or involving direct aPKC phosphoregulation (Atwood & Prehoda 2009) are both correct. Our time-resolved analysis further allowed determining when in the cell cycle the proposed mechanisms operate to establish asymmetric fate determinant localization.

During interphase, Mira localization does not require the actin cortex, but the BH motif and Serine 96 (S96) in particular (**Figure 4A**). Thus, in interphase NBs Mira binds with its BH motif to phospholipids of the PM (Bailey & Prehoda 2015). At the onset of prophase Mira is driven off the PM into the cytoplasm by aPKC since mutating S96 to Alanine prevents clearance and F-actin-independent Mira uniformly binding to the PM persists, which is also true in mitotic *apkc* NBs (**Figure 3A**, **Figure 3 supplement**). In contrast, clearance occurs in LatA or ML-7 treated NBs (**Figure 2A**,**B**). Therefore, aPKC rather than actomyosin dependent processes drive Mira clearance.

In the transition from interphase to mitosis, the NBs cytoskeleton gets remodeled, slowing the diffusion of PM-bound proteins (**Figure 3D**). This stage is also sensitive to low doses of the ROCK inhibitor Y-27632 (**Figure 5B**) and the dynamics of Mira clearance appear altered when LatA treated, cycling NBs enter mitosis (**Movie S3**). Thus, as in the *C.elegans* zygote where actomyosin remodeling contributes to the establishment of polarity (Munro et al. 2004), actomyosin remodeling during prophase might also contribute in NBs to polarize Mira.

After NEB Mira asymmetry requires actomyosin activity (**Figure 2A-D**). Barros et al. (2003) suggested that myosin II is required to exclude Mira from the apical pole pushing Mira it into daughter cells during division. Our results rather suggest that, myosin activity keeps Mira anchored basally (**Figure 2C**). The observation that NEB is a critical time point for basal crescent formation is further in line with the proposition that factors released from the nucleus upon NEB could regulate Mira localization (Zhang et al. 2016). Interestingly, ROCK and MLCK both affect myosin activity (Amano et al. 1996; Ueda et al. 2002; Saitoh et al. 1987; Watanabe et al. 2007) yet the effects of MLCK (ML-7) and ROCK (Y-27632) inhibition on Mira differ (**Figure 2** versus **Figure 5**). In MCDK II cells, Y-27632 and ML-7 treatment has different effects on myosin regulatory light chain phosphorylation (Watanabe et al. 2007). This could result in different effects on myosin activity that may explain the different effects on Mira also in NBs.

Another question raised by our results is whether aPKC contributes to Mira localization following NEB. We attempted to test this by using the ROCK inhibitor Y-27632, also shown to inhibit aPKC (Atwood & Prehoda 2009). High doses (200µM) of Y-27632 applied to cycling NBs recapitulated defective clearance of Mira by aPKC (**Movie S12**), and the same dose applied to already polarized NBs resulted in a partial loss of Mira asymmetry, however after a ~50min delay (**Figure 5E**, **Movie 13**). Thus, aPKC may be required after NEB to reinforce Mira asymmetry by excluding it apically, as suggested (Atwood & Prehoda 2009). Low doses (25µM) of Y-27632, despite not inhibiting aPKC, affect Mira crescent size in cycling NBs (**Figure 5B**). Thus, Y-27632 also inhibits other aPKC-independent mechanisms, making unclear whether the partial loss of Mira asymmetry in already polarized NBs treated with high Y-27632 is solely due to the disruption of a post-NEB aPKC function. Addressing this question in the future will require specific and temporally controlled aPKC inhibition.

When the BH motif is deleted, Mira cannot form crescents (**Figure 4A**). We recently identified that maintenance of Mira crescents also requires interaction of Mira with its cognate mRNA (Ramat et al. 2017). Thus, establishment of Mira crescents and their maintenance are differently controlled. The BH motif might be important to initiate Mira asymmetric localization, while actomyosin dependent processes contribute to establishment and stabilize Mira in a subsequent maintenance step.

Finally, a question of interest is why would Mira localization to the cortex be controlled by different mechanisms in interphase and mitosis. Mira co-localizes with Pros in interphase (**Figure 1 supplement 1**). It is possible that Mira sequesters Pros, helping to prevent its nuclear localization. In this way Pros induced cell cycle exit could be prevented (Choksi et al. 2006; Lai & Doe 2014). Nuclear Pros is developmentally controlled by hedgehog signaling and the temporal transcription factor cascade (Chai et al. 2013; Maurange et al. 2008) and regulated by RanGEF BJ1 (Joy et al. 2014). However, the underpinning cell biology is not clear. Being able to regulate Miras interaction with the PM and at the actomyosin cortex differentially might allow controllable segregation of fate determinants to daughter cells while permitting tuning of nuclear Pros levels through its interaction with Mira in interphase.

## Materials & Methods

### Fly stocks and genetics

Flies were reared on standard corn meal food at 25 degrees. Lines used:

(1) Baz::GFP trap (Buszczak et al. 2007); (2) *w^1118^* (Bloomington); (3) MARCM: hsFlp tubGal4 UASnlsGFP; FRT42B tubGal80/Cyo and FRT82B gal80 (T. Lee & Luo 1999); (4) worniu-Gal4 (Albertson et al. 2004); (5) UAS-Lgl^3A^::GFP (Wirtz-Peitz et al. 2008); (6) UAS-Lgl^3A^ (Betschinger et al. 2003); (7)*w1118, y,w, hsp70-flp; tubP-FRT>cd2>FRT-Gal4, UAS-GFP (Gift from* M. Gho); (8) Mz1061 (Ito et al. 1995); (9) UAS-GFP::Mira (Mollinari et al. 2002). (10) FRT82B *mira^L44^* (Matsuzaki et al. 1998). (10) Df(3R)ora^I9^ (Shen et al. 1997). (11) UAS-aPKC^RNAi^: *P{y[+t7.7] v[+t1.8]=TRiP.HMS01320}attP2* (BL#34332); *(12)* Numb::GFP (Couturier et al. 2013); (13) FRT42B *apkC^k06403^* (Wodarz et al. 2000); (14) UAS-aPKC^ΔN^; (15) P{UASp-sqh.E20E21}3 (BL#64411); (16) P{10xUAS-IVS-mye::tdEos]attP2 (BL #32226); y[1] w[*]; P{y[+t*] w[+mC]=UAS-Lifeact-Ruby}VIE-19A (BL# 35545); (17) aPKC::GFP (Besson et al. 2015); *Source 1 of Mira::mCherry: BAC{mira::mcherry-MS2}* (Ramat et al. 2017). aPKC^RNAi^ clones were generated by heat-shocking larvae of the genotype *y,w, hsp70-flp; tubP-FRT>cd2>FRT-Gal4, UAS-GFP; P{y[+t7.7] v[+t1.8]=TRiP.HMS01320}attP2.* Heat shocks were performed 24hph and 48hph for 1 hour at 37°C. MARCM clones were generated by heat shocking L1 larvae for 2h at 37°C.

Generation of Mira alleles: Source 2 of Mira::mCherry: mira^mCherry^; mira^ΔBHrncherry^ (Ramat et al. 2017), mira^S96A-mCherry^ and mira^S96D-mCherry^ are derived from mira^KO^ (Ramat et al. 2017). mira^mCherry^ was generated by inserting a modified wt genomic locus in which mCherry was fused to the C-terminus following a GSAGS linker into *mira*^ko^. For mira^S96D-mCherry^: TCG (Serine96) was changed to GAC (aspartic acid). For mira^S96A-mCherry^: TCG was replaced with GCG (alanine). CH322-11-P04 was the source for the *mira* sequences cloned using Gibson assembly into the RIV white vector (Baena-Lopez et al. 2013) that was injected using the attP site in mira^KO^ as landing site. *BAC{mira::mcherry-MS2}* (Ramat et al. 2017) **see Figure 1 supplement 2**). While *mira^mCherry^* behaves similarily to *BAC{mira::mcherry-MS2}* and rescues embryonic lethality, *mira^ΔBHmCherry^, mira^S96A-mCherry^* and *mira^S96D-mCherry^* are homozygous lethal.

Live imaging: Live imaging was performed as described (Pampalona et al. 2015). Briefly, brains were dissected in collagenase buffer and incubated in collagenase for 20 minutes. Brains were transferred to a drop of fibrinogen (0.2mgml^-^1, Sigma f-3879) dissolved in Schneider’s medium (SLS-04-351Q) on a 25mm Glass bottom dish (WPI). Brains were manually dissociated with needles before the fibrinogen was clotted by addition of thrombin (100Uml^-1^, Sigma T7513). Schneider’s medium supplemented with FCS, Fly serum and Insulin was then added. A 3-4µm slice at the center of the neuroblasts was then imaged every 30-90s using a 100x OIL objective NA1.45 on a spinning disk confocal microscope. Data was processed and analyzed using FIJI (Schindelin et al. 2012). For nuclear volume measurements Imaris was used. All other drugs were added to the media either prior or during imaging: ML-7 (Sigma, I2764, dissolved in water), Y-27632 (Abcam, Ab120129, dissolved in water). Drugs were washed out by media replacement, in the polarity reconstitution assay colcemid concentrations were kept constant throughout the experiments. FRAP experiments were carried out on a Leica SP8 confocal using a 63x NA1.2 APO water immersion objective. To estimate t_1/2_ for the recovery curves we used published curve fitting methods (Rapsomaniki et al. 2012).

### Immunohistochemistry

#### Primary cell culture

Brains were dissected in collagenase buffer and incubated for 20 minutes in collagenase, as for live imaging. Brains were then transferred into supplemented Schneider’s medium and manually dissociated by pipetting up and down. Cells were pipetted onto a poly-lysine coated 25mm glass bottomed dish and left to adhere for 40 minutes. Schneider’s was then replaced with 4% Formaldehyde (Sigma) in PBS and cells were fixed for 10 minutes. Cells were permeabilized with 0.1% PBS-Triton for 10 minutes. Cells were then washed with PBS 3x 10minutes before antibody staining overnight at 4°C. All antibodies were dissolved in PBS-1%Tween. *Whole mount brains:* Brains were fixed in 4% Formaldehyde (Sigma) for 20 minutes at room temperature. Primary antibodies: Rabbit anti-Miranda (1:200, gift from C. Gonzalez), Mouse anti-GFP (1:400, Abcam). Rabbit anti-Brat (1:200, a gift from J. Knoblich). Guinea Pig anti-Dpn (1:500 a gift from J. Skeath). Mouse anti-Pros (1:40, DSHB). To stain F-Actin we used Alexa Fluor 488 or 561 coupled Phalloidin (Molecular Probes, 5:200) for 20 minutes at room temperature. Secondary antibodies (all from life technologies and raised in donkey: Anti-Rabbit Alexa-594, Anti-Mouse Alexa-488, Anti-Rabbit Alexa-647, Anti-Guinea Pig Alexa-647. Microscopy was performed using a Leica-SP8 CLSM (60x Water objective, 1.2) and images were processed using FIJI.

In all cases the sample size *n* provided reflects all samples collected for one experimental condition. Experimental conditions were repeated at least twice to account for technical and biological variation.

## Acknowledgements

We thank C. Doe, J. Knoblich, F. Schweisguth, D. StJohnston, F. Matsuzaki, C. Gonzalez, J. Skeath, A. Wodarz, M. Gho and the Kyoto and Bloomington stock centers for reagents and/or protocols. We thank A. Müller, C. Weijer and M. Gonzalez-Gaitan for critical reading. M.R.H is supported by an MRC PhD studentship. We thank inDroso (http://www.indroso.com) for the generation of *mira^attPko^* by CrispR and the Dundee imaging facility for excellent support. Work in J.J.’s laboratory is supported by Wellcome and the Royal Society Sir Henry Dale fellowship 100031Z/12/Z. M.R.H is supported by an MRC studentship funded by these grants: G1000386/1, MR/J50046X/1, MR/K500896/1, MR/K501384/1. The tissue imaging facility is supported by the grant WT101468 from Wellcome.

## Figure supplements

**Figure 1 supplement 1.**
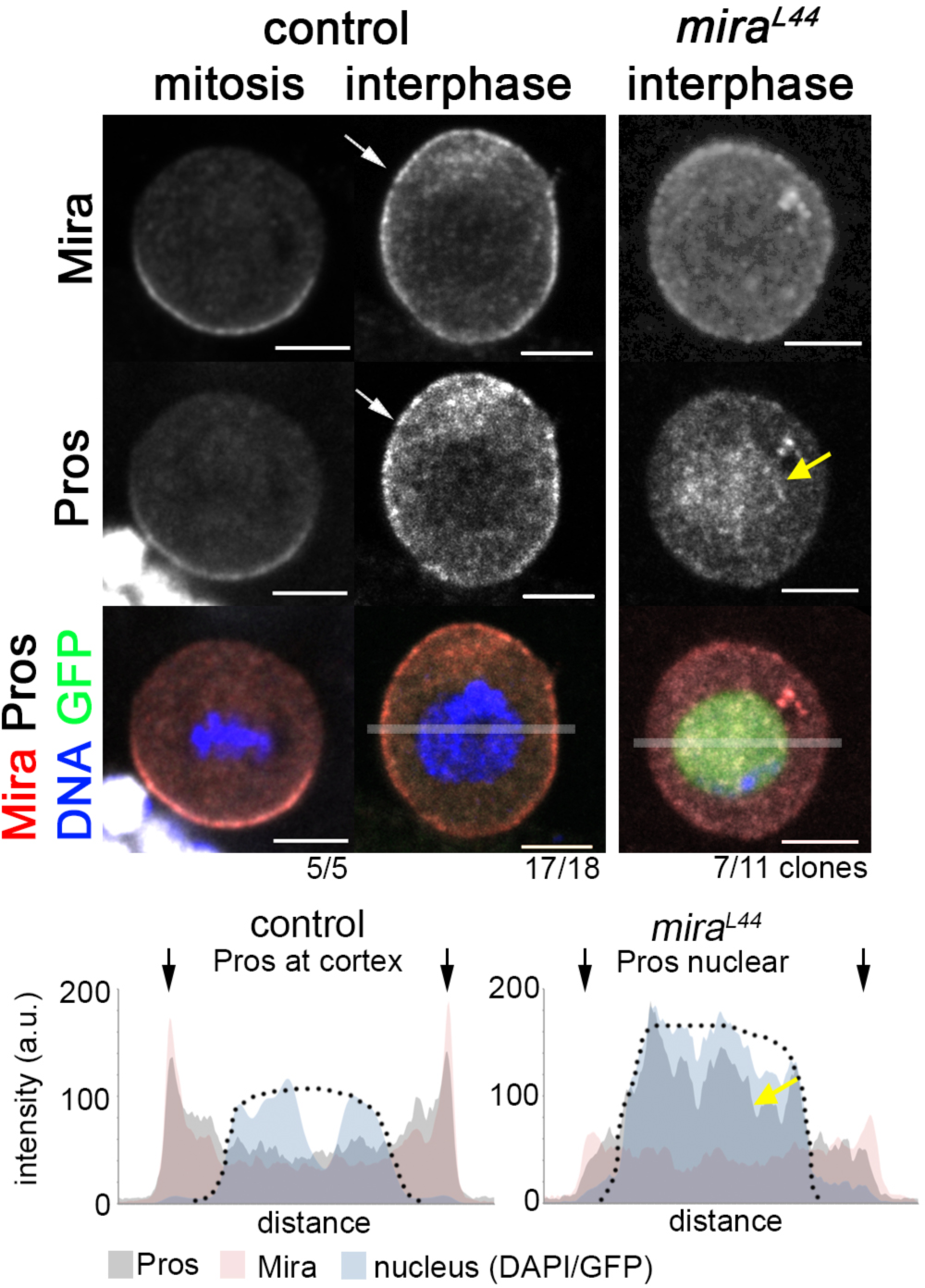
Uniform cortical Prospero depends on Miranda in interphase larval NBs. In w^1118^ brains, Mira and Pros form basal crescents in mitosis and both are cortical in interphase (arrow). In an interphase *mira*^144^ NB (MARCM clone, GFP^+^) cortical Mira and Pros are strongly reduced and Pros accumulates in the nucleus (yellow arrow). Transparent bars in merge interphase (control) and *mira*^144^ indicate area used for plot profiles shown below. Pros and Mira are at the cortex (arrows) in the control and Pros is enriched in the nucleus in the mutant (yellow arrow). Arrows: outline of cell. Dotted line: nucleus (based on DAPI, control and GFP (MARCM clone). Scale bar: 10µm.

**Figure 1 supplement 2.**
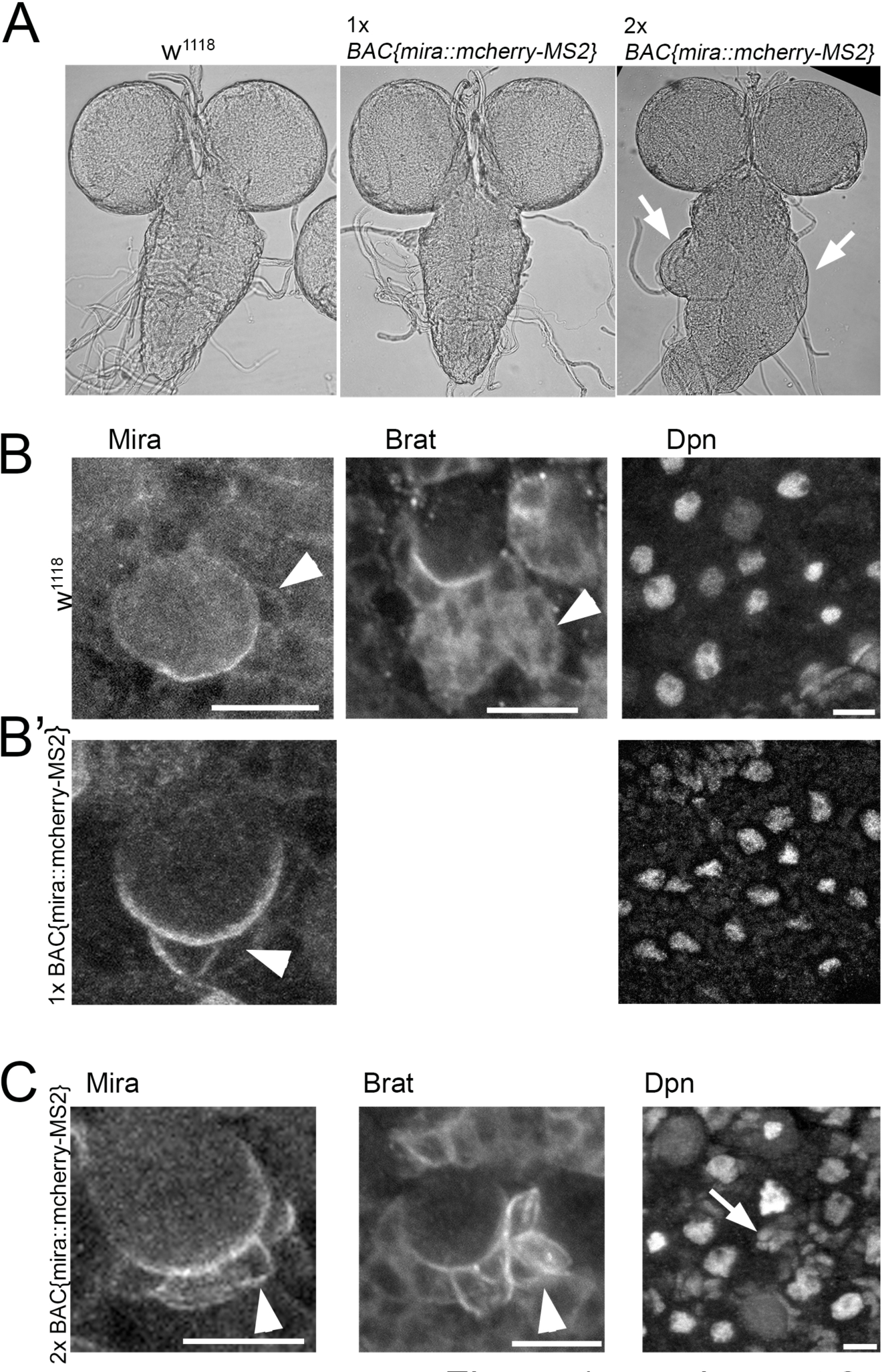
***BAC{mira::mcherry-MS2}*** rescues embryonic lethality of the loss of function allele mira^L44^ over the deficiency DF(3R)ora^I9^. However, animals die during puparium formation, when *BAC{mira::mcherry-MS2}* is the only source of Mira. (**A**) Brightfield images of fixed whole mount brain preparations. *W^1118^* (control, n=5) a *BAC{mira::mcherry-MS2}* brain over a wild type chromosome (1x *BAC{mira::mcherry-MS2}, n=12)* and a brain from *BAC{mira::mcherry-MS2}* Df(3R)ora^I9^ over a unrecombined *mira^BACmcherry^* chromosome (2x *BAC{mira::mcherry-MS2}, n=12).* 2x *BAC{mira::mcherry-MS2}* animals die as pharates. The ventral ganglion (VG) of these brains is frequently overgrown (arrows). Similar effects are seen with CrispR generated, homozygous *mira::mCherry::HA* larvae (not shown). (**B**) In fixed ***w****^1118^* brains Mira as well as its cargo Brat are diffuse in the cytoplasm of NB daughter cells and Deadpan (Dpn) staining is restricted to NB nuclei. (**B’**) *BAC{mira::mcherry-MS2}brains* are not overgrown, Mira is sometimes more stable at the cortex in a daughter cell (arrowhead), but Dpn is normal. (**C**) In 2x *BAC{mira::mcherry-MS2}* animals, Mira is strongly cortical in several NB daughter cells and so is Brat (arrowheads). Dpn is no longer restricted to NB nuclei but frequently found in clusters of smaller nuclei close to NBs suggesting that Mira is stabilized at the cortex and fails to release its cargo, inducing fate changes. mCherry is fused to the C-terminus of Mira which was shown to be required for cargo release (Fuerstenberg et al. 1998; Matsuzaki et al. 1998). Scale bars: 10µm.

**Figure 1 supplement 3.**
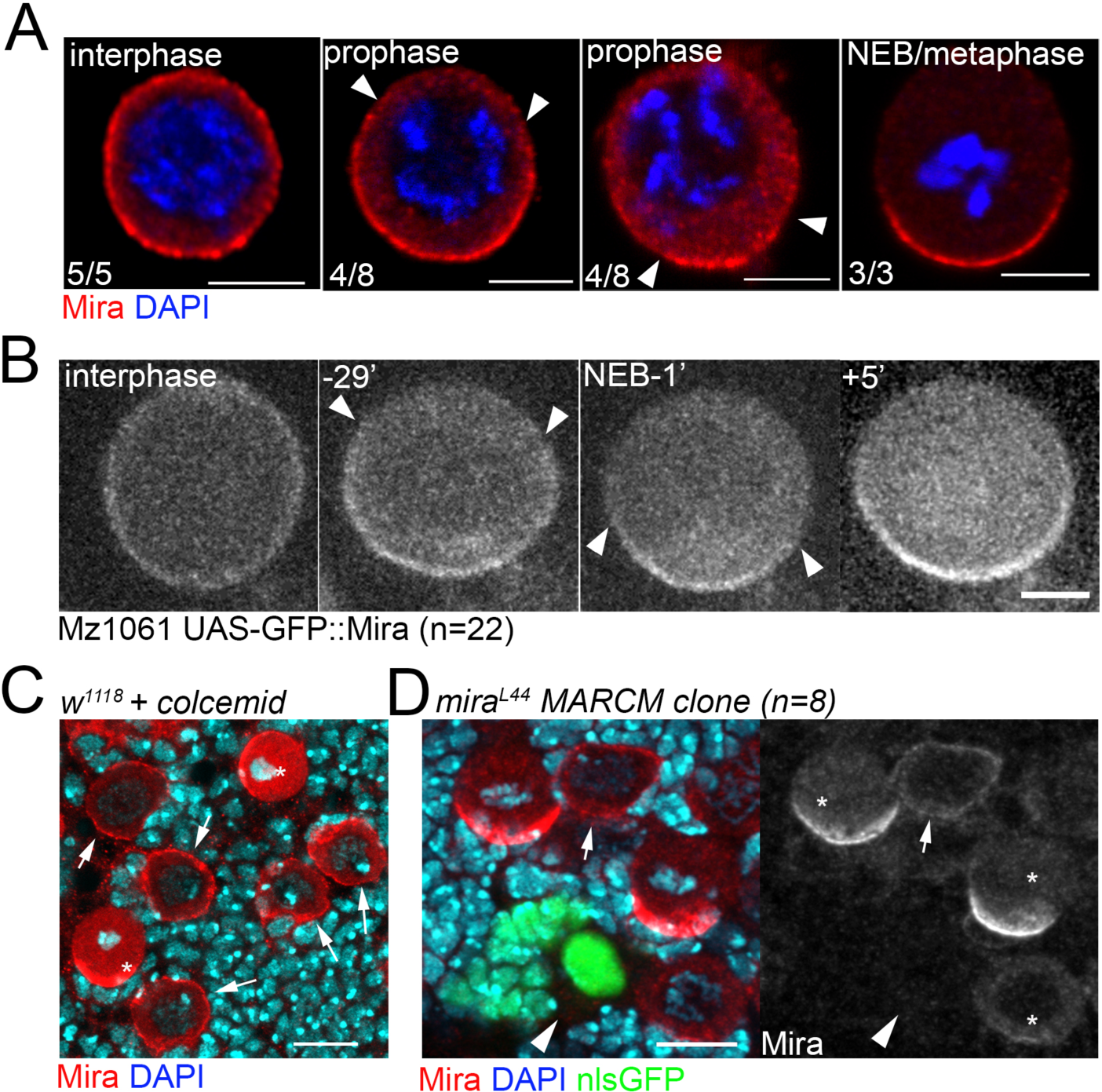
Cortical Mira can be detected by antibody staining, in UAS-GFP-Mira overexpressing NBs and upon colcemid treatment, but not in interphase mira^L44^ loss of function clones. (**A**) Antibody staining against Mira performed on different fixed isolated NBs in primary cell culture. In this assay Mira (red) is cortical in an interphase NB (judged by DAPI, blue). In prophase NBs, differently sized Mira “crescents” can be detected the ends of which are labeled by arrowheads. In a metaphase NBs Mira forms a crescent that appears larger than some of those seen in prophase NBs. (**B**) A living NB in primary cell culture expressing Mira : :GFP driven by Mz1061. GFP signal is at the cortex in interphase, 29 min prior to NEB, GFP signal becomes cleared apically (arrowheads) until most of the cortex is cleared 1min prior to NEB. 5 min after NEB a robust, larger crescent has formed. (**C**) Control brain treated with 50µM colcemid for 30min and stained for Mira. Over condensed chromatin in mitotic NBs demonstrates the effect of colcemid yet in all interphase NBs Mira remains at the cortex. (**D**) Fixed brain containing an interphase NB MARCM *mira^L44^* clone surrounded by control NBs. In the clone cortical Mira signal is gone (arrowhead) while present in a control interphase NB (arrow). Asterisks: mitotic NBs. Scale bar 10µm.

**Figure 2 supplement.**
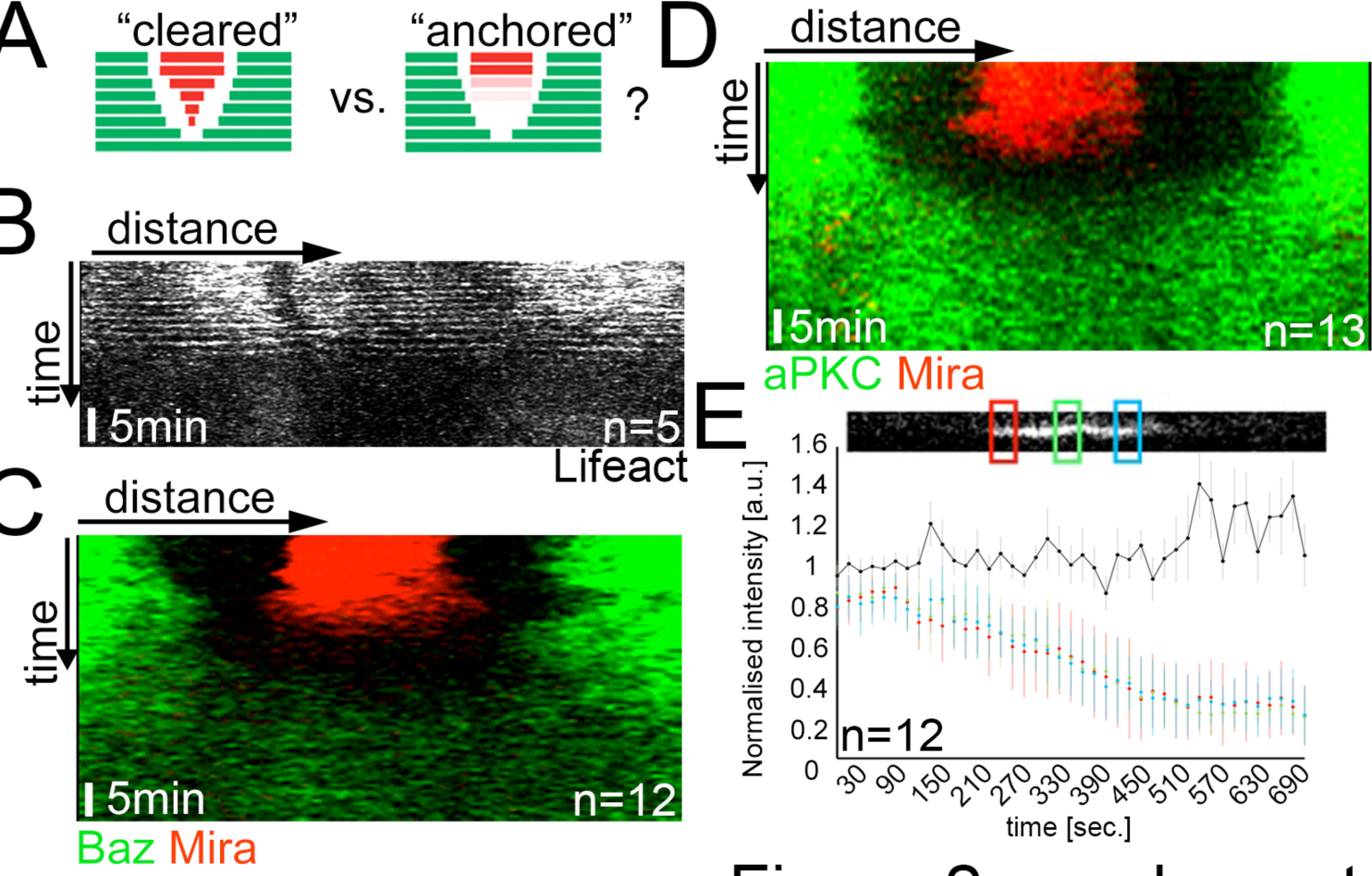
Mira falls homogenously off the cortex upon LatA treatment, which is not driven by aPKC cortical displacement. (**A**) Schematic depicting the expected kymograph profile for *clearing* versus *anchoring.* (**B-D**) Kymographs of colcemid arrested NBs expressing Lifeact-Ruby, Baz::GFP and Mira::mCherry or aPKC::GFP and Mira::mCherry (related to **Movie S4**) upon the addition of 5µM LatA. The equatorial perimeter of the NB was straightened out for each time point. Time scale bar: 5min. (**E**) Plot of Mira decline upon LatA treatment of a colcemid arrested NB using 3 ROIs (positions as indicated). Black line shows ratio ROI3/ROI2 over time, which remains constant. Error bars: standard deviation. *BAC{mira::mcherry-MS2}* was the source of Mira::mCherry. Scale bar: 10µm.

**Figure 3 supplement.**
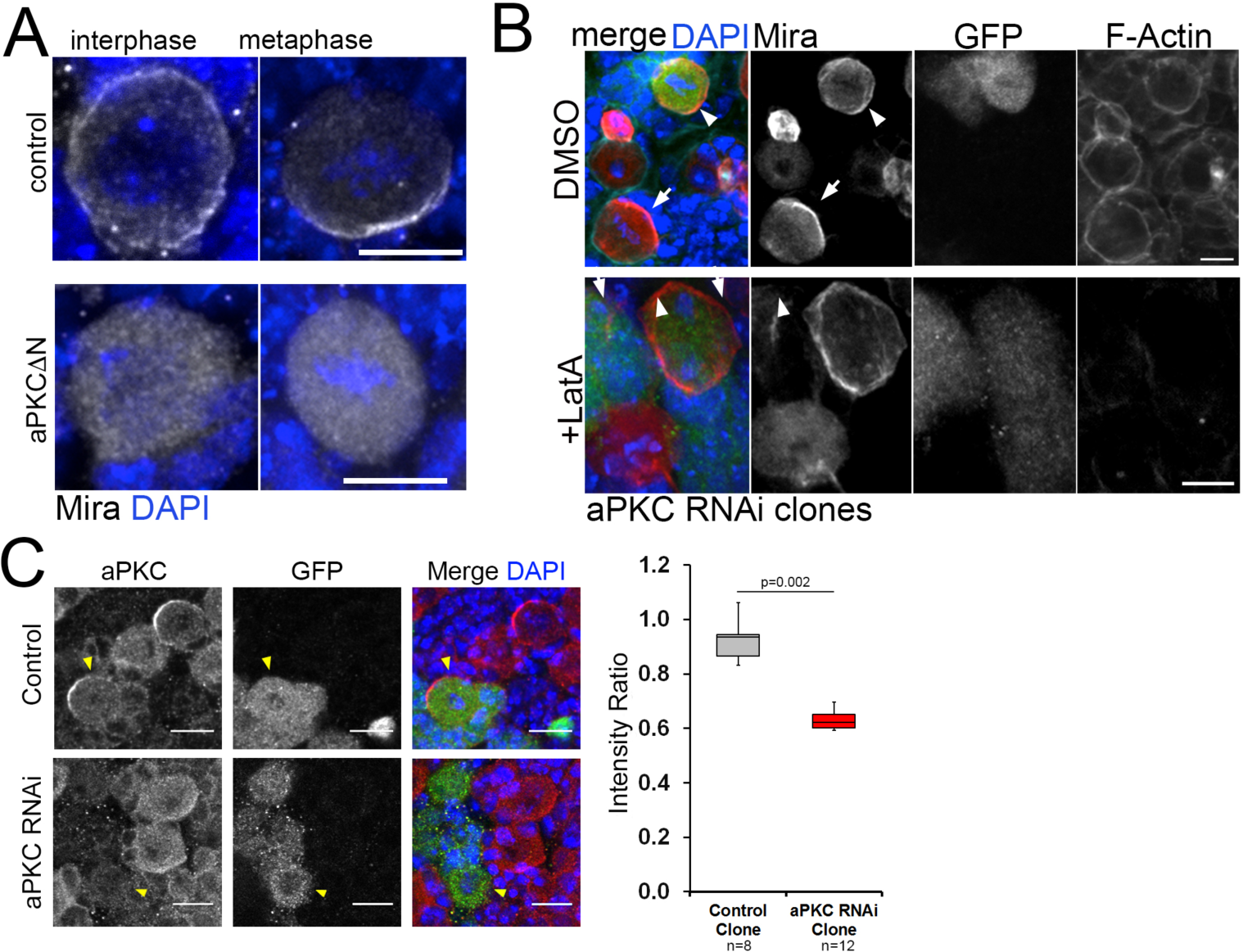
Effects of aPKC manipulation on Mira localization under different conditions. (**A**) Mira antibody staining on whole mount brains in a control NBs and in a NBs that expresses aPKC^ΔN^ by worniuGal4. Control: Mira is cortical in interphase (91% cortex, 9% cytopl., n=53), forms a crescent in metaphase (93% crescent, 7% cytopl., n=15). aPKC^Δ N^: Mira is cytoplasmic in interphase (85% cytoplasm, 15% cortex, n=40) and metaphase (89% cytopl., 11% crescent, n=15). (**B**) aPKC RNAi expressing flip out clones (GFP positive, arrowheads) and GFP negative control NBs (arrows) treated with DMSO or 5µM LatA and stained with an antibody against Mira and with Phalloidin to label F-Actin. In GFP negative mitotic control NBs, Mira is in a crescent (DMSO) or cytoplasmic (LatA). Mira is cortical in DMSO as well as LatA treated mitotic aPKC RNAi NBs (100%, n=5, arrowheads). (**C**) aPKC is efficiently knocked down by RNAi in flip out clones. aPKC RNAi NBs have significantly less aPKC. Scale bar: 10µm.

**Figure 5 supplement.**
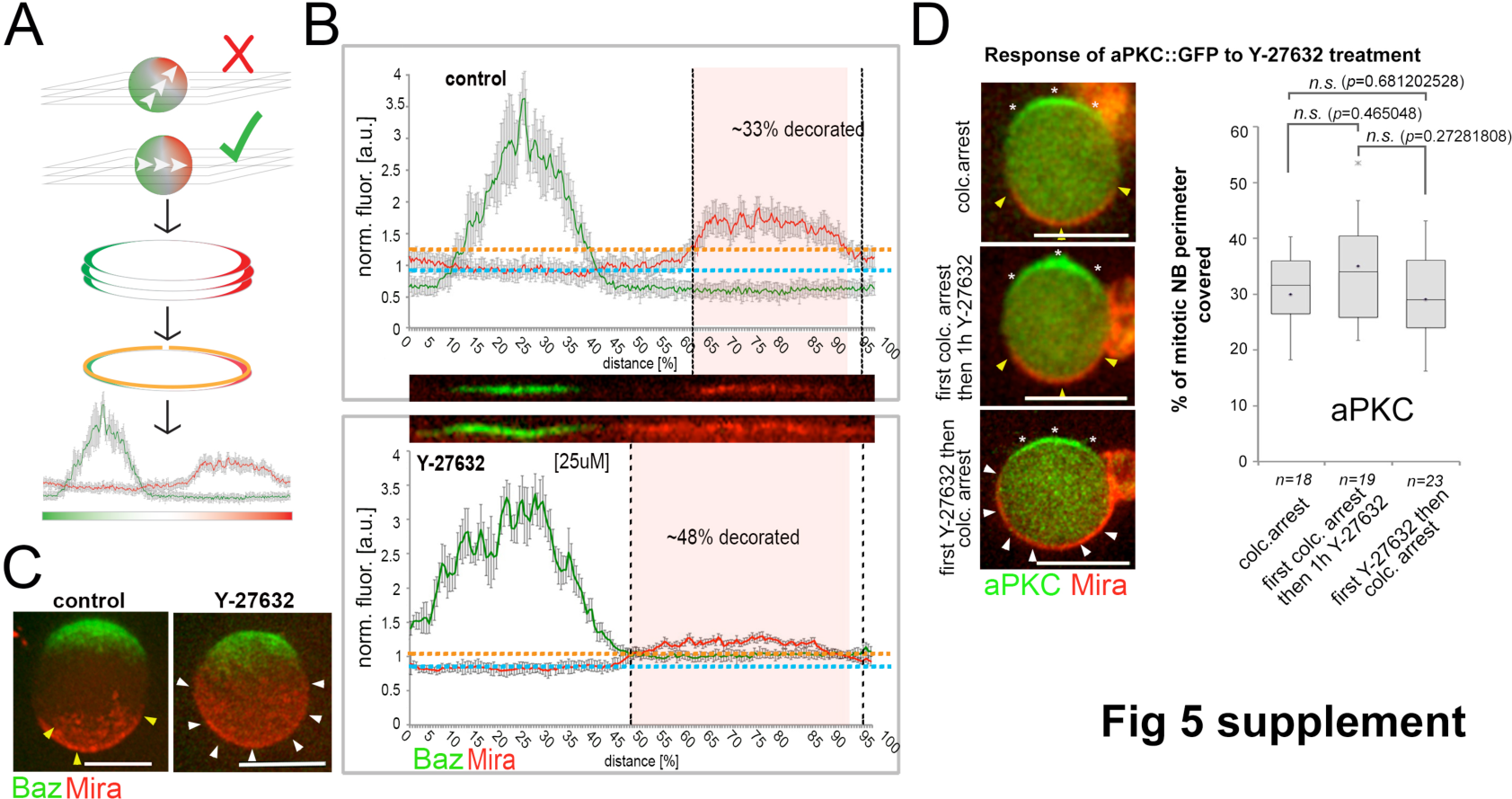
Standard used to quantify Mira crescent size. (**A**) Schematic of workflow. NBs that have a polarity axis parallel to the imaging plane are selected. 3-5 optical planes are collected covering 2-3µm of the equator of the NB. (**B**) Fluorescence is normalized against the cytoplasmic background and straightened line plots are derived from each section. The average background and the average standard deviation is determined. Signal: > avg. background plus two times the average standard deviation. (**C**) 3D projections of z-sections covering the entire NBs, ctrl vs. a NB that polarized in the presence of 25µM Y-27632. (**D**) Quantification of aPKC::GFP crescent size under the indicated conditions (unpaired ttest). Asterisks: aPKC; yellow arrowheads: normal sized crescents; white arrowheads: enlarged Mira crescents. *BAC{mira::mcherry-MS2}* was the source of Mira::mCherry. Scale bar: 15µm.

## Movie captions

### Movie S1

Interphase cortical Miranda is removed at the onset of mitosis. Spinning disc confocal image of a neuroblast expressing Baz::GFP (red) and Mira::mCherry (green). For this and all subsequent videos maximum projection after a 3D Gaussian blur (FIJI, radius 8/.8/1) of 7 consecutive equatorial planes taken at 0.4μm spacing is shown. Z-stacks taken every minute. Time stamp: hh:mm.

### Movie S2

Interphase cortical Miranda is removed at the onset of mitosis. Spinning disc confocal image of a neuroblast expressing aPKC::GFP (green) and Mira::mCherry (red). Z-stacks taken every minute. Time stamp: hh:mm.

### Movie S3

Interphase cortical Miranda is actin independent. Spinning disc confocal image of a neuroblast expressing Baz::GFP (red) and Mira::mCherry (green) showing a control division before 1μM LatA was added. Z-stacks taken ~every minute. Z-stacks taken every minute. Time stamp: hh:mm.

### Movie S4

Colcemid arrested NBs expressing aPKC::GFP and Mira::mCherry that were treated with 5µM LatA at the beginning of the recording at 16sec intervals. The cortex was straightened out and split at the apical pole such that aPKC::GFP appears right and left and Mira in the centre. Fluorescence profiles shown below. Note that Mira falls off homogenously from the cortex and becomes cytoplasmic at 5:36 (red arrowhead), while the detectable borders of cortical aPKC (green arrowheads) have not yet changed. Only from 7:12 onward aPKC rise above cytoplasmic levels where Mira was localized. Time stamp: mm:ss.

### Movie S5

Myosin inhibition reversibly affects basal Mira anchoring in a polarized neuroblast. A colcemid arrested NB expressing Baz::GFP (green) and Mira::mCherry (red) in primary cell culture was treated with 20µM ML-7 which as washout when indicated. Left panel Baz::GFP, middle panel Mira::mCherry, right panel merge. Z-stacks taken every minute. Time stamp: mm:ss.

### Movie S6

The effect of ML-7 on cortical Mira localization in mitosis can be delayed by overexpressing Sqh^EE^. Mira::mCherry NBs (ctrl) and Mira::mCherry NBs co-expressing Sqh^EE^ (rescue) were co cultured in neighboring clots in the same dish and the effect of ML-7 on cortical Mira recorded. Z-stacks taken every two minutes. Time stamp: hh:mm.

### Movie S7

Miranda remains at the cortex throughout the cell cycle in *apkC^k04603^* mutant NBs. Spinning disc confocal image of an apkc^k04603^ mutant NB, labeled with nlsGFP (green) expressing Mira::mCherry (white). Z-stacks taken every minute. Time stamp: hh:mm.

### Movie S8

Control mira^mCherry^ allele generated by CrispR/Cas9. Mira localizes to the interphase cortex, from where it is cleared before NEB. Then Mira re-localizes to a larger crescent. Therefore this allele and Mira::mCherry (BAC rescue) are undistinguishable in terms of Mira dynamics. This control further shows that the MS2 binding site in the BAC rescue construct does not interfere with Mira cortical dynamics. Time stamp: hh:mm.

### Movie S9

Phosphomutant S96A allele of Mira tagged with mCherry at the C-terminus. Mira localizes uniformly to the interphase cortex. Shortly before NEB, S96A is apically enriched, before being uniformly cortical after NEB and during division. Time stamp: hh:mm.

### Movie S10

Phosphomimetic S96D allele of Mira tagged with mCherry at the C-terminus. S96D localizes to microtubules in interphase, but is asymmetric in mitosis in mitosis. Note the signal resembling subcortical microtubules in interphase converging at the apical pole. After NEB a basal crescent is detectable. At 115:30 a z-stack spanning the entire NB was collected and the maximum projection is frozen. After this 50µM colcemid was added to reveal if Mira^S96D^::mCherry binds to the cortex. Next frozen frame: similar stack after 30min in colc. Next frozen frame: 50 in colcemid – no cortical signal is detectable. Last frozen frame 65min in colcemid. Time stamp: mm:ss. Scale 15µm.

### Movie S11

Mira requires its BH motif for interphase cortical localization (see main text) and basal localization in mitosis. The BH motif in Mira has been deleted by gene editing and this Mira mutant tagged with mCherry at the C-terminus (*mira^ΔBHmCherry^*). Mira^ΔBH^::mCherry when homozygous is found on the interphase microtubule network and in the cytoplasm during mitosis. Time stamp: hh:mm.

### Movie S12

200µM Y-27632 induced uniform cortical Mira in mitosis localizes independently of an intact actin network. A Baz::GFP and Mira::mCherry expressing NBs was cultured in the presence of 200µM Y-27632, arrested with colcemid. 5µM LatA was added after the first frame of the movie. LatA induces loss of Baz asymmetry, yet Mira remains cortical. Z-stacks shown. Z stacks taken every 2min. Time stamp: hh:mm.

### Movie S13

Colcemid arrested NBs expressing Baz::GFP and Mira::mCherry, treated with 200μM Y-27632. Mira starts to become visible ~36min after Y-27632 addition in this example, but remaines asymmetrically distributed, until LatA is added. Z-stacks shown. Z stacks taken every 2min. Time stamp: mm:ss.

## Author contribution

M.H., A.R., N.L. and J.J. designed and carried out experiments and interpreted the data and M.H., N.L. and J.J. wrote the manuscript that was agreed upon by all authors.

